# PyDecNef: An open-source framework for fMRI-based decoded neurofeedback

**DOI:** 10.1101/2023.10.02.560503

**Authors:** Pedro Margolles, Najemeddine Abdennour, Ning Mei, Patxi Elosegi, David Soto

## Abstract

Real time fMRI research has suffered from inaccessible analysis pipelines, hindering collaboration and reproducibility. Here we present PyDecNef, a Python-based platform designed to advance real-time fMRI analysis and fuel exploration of close-loop neuroimaging for cognitive neuroscience studies. Creating a real-time fMRI analysis pipeline from scratch poses formidable technical challenges, involving data transfer, experimental software, and machine learning classifier preparation. Existing tools like FRIEND, Brain-Voyant, and OpenNFT demand expensive licenses or rely on proprietary software, impeding accessibility and customizability. PyDecNef offers a solution: a transparent, versatile, and open workflow for real-time fMRI decoding protocols. This open-source platform simplifies decoder construction, real-time preprocessing, decoding, and feedback signal generation. It also supports co-adaptive decoders that update in real time based on decoded neurofeedback performance, enabling researchers to conduct more precise and efficient DecNef experiments. Moreover, its openness promotes collaboration, enhancing research quality, replicability, and impact. With PyDecNef, the path to advancing DecNef studies becomes more accessible and collaborative. PyDecNef resources for real-time fMRI analysis can be found at https://github.com/pydecnef/PydecNef.

## Introduction

One of the main challenges for the development of decoded neurofeedback (DecNef) studies is setting up a real-time fMRI processing framework. Until now, DecNef studies employed analysis pipelines that were not openly available to the scientific community. We introduce PyDecNef, a Python-based, flexible, fully open platform for real-time fMRI analysis that we make openly available to advance research in this area, foster collaborative efforts and reproducibility, and promote the emergence of new research lines in DecNef studies. At the same time, DecNef research faces a second, equally important challenge: a substantial proportion of participants struggle to learn how to reliably induce the target brain pattern during training, in part because the decoder trained on one dataset (e.g. perceptual runs) may not generalise optimally to the neural activity generated during neurofeedback sessions. PyDecNef was designed to tackle both obstacles by providing not only an accessible real-time fMRI pipeline, but also dedicated tools to improve decoder calibration and adaptability so that feedback more effectively supports participants’ learning.

In parallel with these technical barriers, neurofeedback and DecNef studies often report modest or absent learning curves for many individuals. This difficulty is likely exacerbated by differences in data distributions between the training data used to construct the decoder and the data generated during neurofeedback test sessions, as well as by neural and non-neural noise and non-stationarities in fMRI signals. In our initial development of PyDecNef, we therefore focused on enhancing the precision and generalizability of the decoder in a real-time fMRI context, by implementing a customised classification pipeline that yields more graded and better calibrated probability estimates across perceptual, imagery and neurofeedback data. We have also extended this approach by incorporating a co-adaptive training module: rather than only asking participants to learn how to induce the target state, the decoder itself can also adapt online to high-confidence decoded neurofeedback examples, thereby reducing the mismatch between training and test distributions and facilitating participants’ ability to reach the target state during DecNef.

Building from scratch a fMRI real-time analysis pipeline thus entails a substantial technical challenge for any researcher that would like to enter the field of DecNef. Multiple components are involved, namely: managing data transfer between the MRI scanner, the real-time analysis PC, and the experimental software that controls the stimuli, task and neurofeedback signals given to the participant. A key step in this process involves the preparation of the machine learning classifier for real-time application. Here it is critical not only to enhance the precision and generalizability of the classifier, but also to ensure that the framework is flexible enough to support both static and co-adaptive decoding strategies, and to monitor decoder stability across sessions. When we started to develop PyDecNef, there were a few software packages to perform decoded neurofeedback studies, namely, the FRIEND toolbox (Sato et al., 2018), Brain-Voyant (Cohen, Malach, Koppel, & Friedman, 2018), Turbo-BrainVoyager (Goebel, 2012), BrainIAK (Kumar et al., 2019) and OpenNFT (Koush et al., 2017). However, some of these packages are commercial, requiring the acquisition of an expensive license or relying on proprietary and closed-source MATLAB software, while others lack customizability or clear documentation.

We believe that a truly open, simple but flexible and customizable workflow for real-time fMRI analysis and decoded neurofeedback would be extremely valuable for any research group thinking about starting to run their own DecNef experiments, saving them time, money and also a few headaches along the way. Moreover, an open pipeline for DecNef would facilitate collaboration across many laboratories, enhancing the quality, the replicability and the impact of DecNef research. With these motivations in mind, we decided to develop and release our open-source fMRI-based DecNef pipeline, and to progressively enrich it with methods that directly address participants’ difficulty in inducing the target state, such as improved decoder calibration and co-adaptive training procedures.

PyDecNef is available at https://github.com/pydecnef/PydecNef. It provides an open and easy-to-use framework for performing real-time fMRI decoded neurofeedback studies using Python 3.11.7 or later, which we recommend for optimal compatibility. The scripts are also backward compatible with the previous release, which supports Python 3.6 and above. The library includes documented scripts for decoder construction and for real-time fMRI preprocessing, online decoding, feedback signalling, and (optionally) co-adaptive updating of the decoder. Overall, the PyDecNef framework provides the researcher with a number of benefits over other real-time fMRI analysis software, namely:

- *Customizable*: pyDecNef code frontend was written in Python 3.11. Python stands out for being an open, powerful, flexible, user-friendly and efficient high-level language. This feature in combination with the modular and object-oriented programming approach used for the development of the pipeline allows for easy customization of DecNef paradigms without the need of deep coding skills.
- *Adaptable*: PyDecNef can be easily adapted for the execution of other type of neurofeedback studies that involve self-regulation of brain activity levels within a target region of interest (e.g. Scharnowski et al., 2012) or to further develop other closed-loop fMRI protocols, for instance applying Bayesian Optimization to delineate the key stimulus parameter space that is most suitable to maximize a relevant neural pattern in the brain (Lorenz et al., 2016).
- *Integral*: Our framework provides a complete approach for the development of DecNef experiments, including tools for processing and decoding fMRI data in real-time and raw example data and resources for different decoder construction, DecNef approaches (i..e. static vs dynamic), off-line analyses, and experimental software for stimulus and feedback presentation.
- *Simple*: pyDecNef framework is easy to use and understand, whether you are a neuroimage novice or an expert. All scripts are well documented.
- *Fast*: While Python is known to be a highly-interpretable and easy to use programming language, it does not stand out for its processing speed. Computational efficiency is an essential requirement in real-time MRI settings, specially in DecNef paradigms where feedback given to participants need to be as close as possible to target the neural pattern of interest. By means of the Nipype library (Gorgolewski et al. 2011), pyDecNef incorporates functionalities from computationally efficient neuroimage software written in C language (i.e. Afni). This makes pyDecNef both flexible and fast.
- *Fully open-access*: Last but not least pyDecNef is an open-access, collaborative and free to use, which will improve henceforth owing to the efforts of the scientific community. Technical support and consulting is also freely given to any researcher intending to run a DecNef experiment using our pipeline.

### Installation of the library

To set up the necessary infrastructure, an Ethernet Local Area Network (LAN) connecting is needed three computers:

#### MRI Scanner Host Computer

The pyDecNef framework is compatible with various types of MRI scanners. However, for the smooth integration with FIRMM software, necessary for transferring volumes from the MRI scanner host computer to the real time analysis computer (i.e. the server), it is recommended to use Siemens or GE scanner models. The host computer is used for managing scanning sessions and transferring the data to the server computer.

The MRI scanner host computer should be configured to copy functional DICOM files to a designated folder on the server computer in real-time. The server computer runs real-time processing scripts, including a watcher class responsible for monitoring the folder and initiating volume preprocessing as soon as a new volume is written to it. To achieve this, Siemens Trio and Prisma scanner host computers can use the “ideacmdtool” program in conjunction with the FIRMM software, while GE scanners can utilize the rsync protocol.

<u>Using ideacmdtool + FIRMM with SIEMENS scanners</u>
<u>Using rsync + FIRMM with GE scanners</u>
<u>FIRMM installation</u>

#### Server Computer

The server computer is responsible for running real-time volume preprocessing and decoding scripts.

#### Client Computer

This computer runs the Python-based software for presenting stimuli and feedback to the participant. Example scripts are included to run in Opensesame (Mathôt, Schreij & Theeuwes, 2012)

<u>Opensesame installation</u>

Figure 1 below illustrates the set-up.

**Figure 1.**
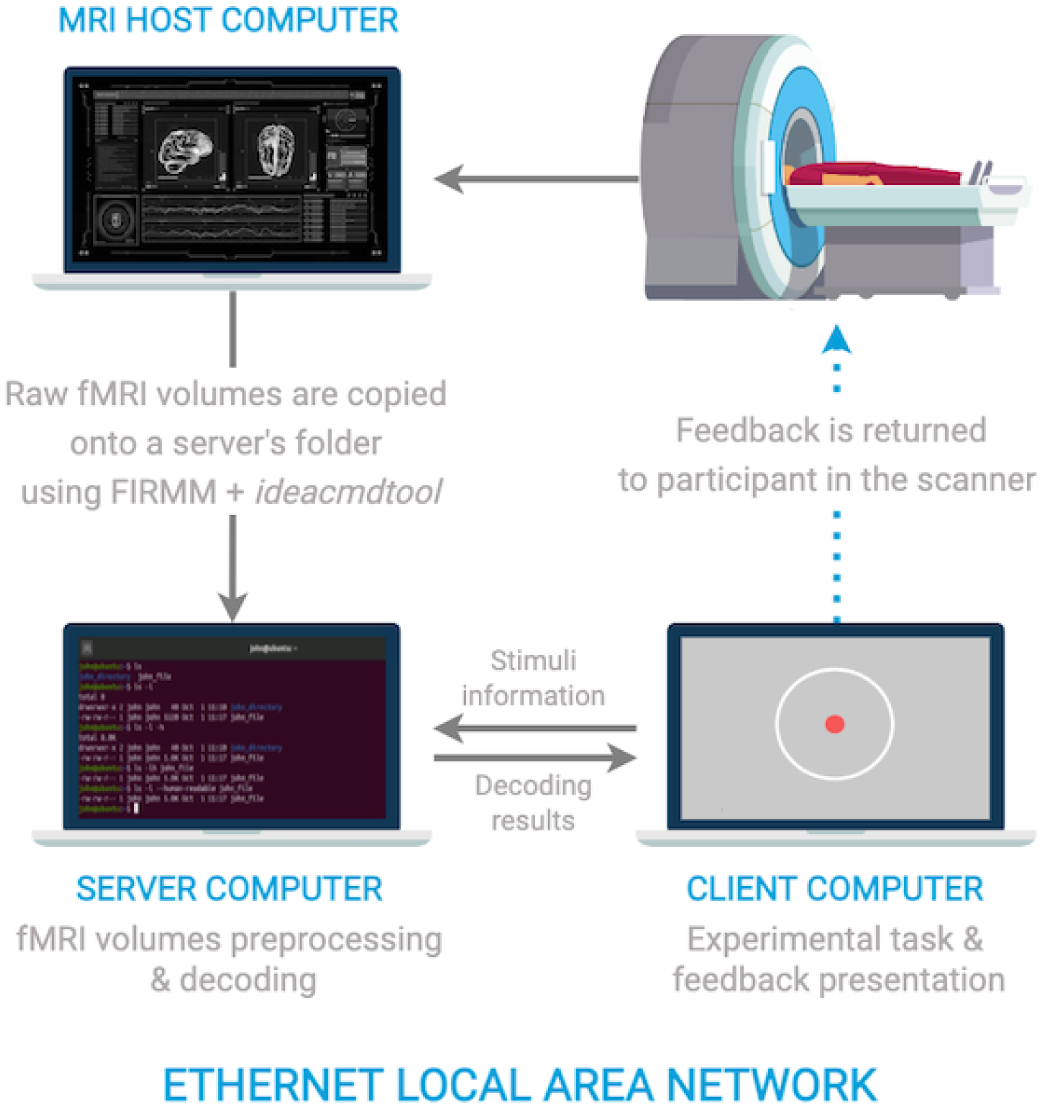
Illustration of the setup of the three computers needed to perform real-time fMRI.

PyDecNef follows the structure shown in Figure 2, which relies on two main computers: a server, which handles real-time fMRI data processing, and a client, which manages stimulus presentation and feedback delivery.

**Figure 2.**
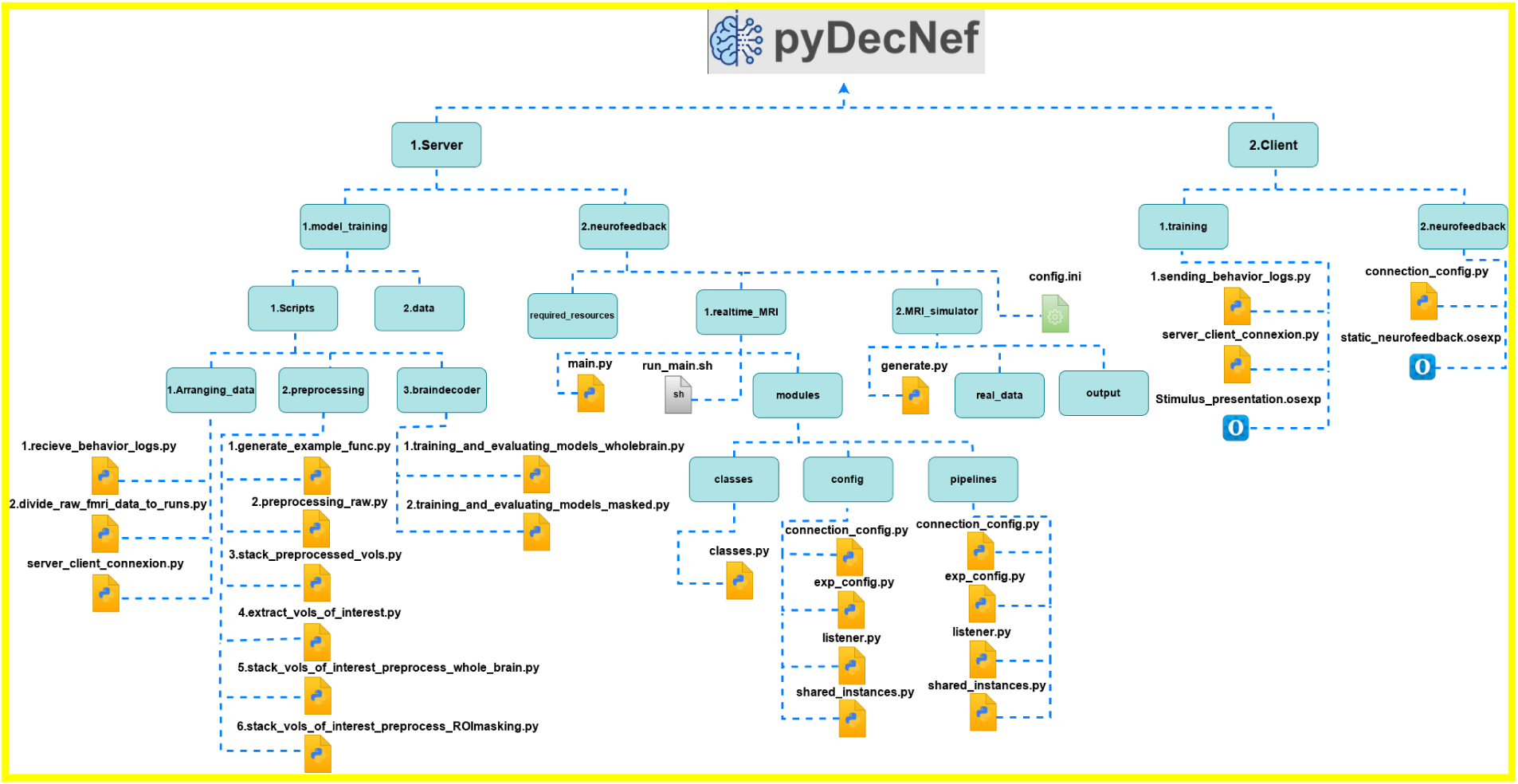
Illustration of the file tree of the PyDecNef package.

**Figure 2.**
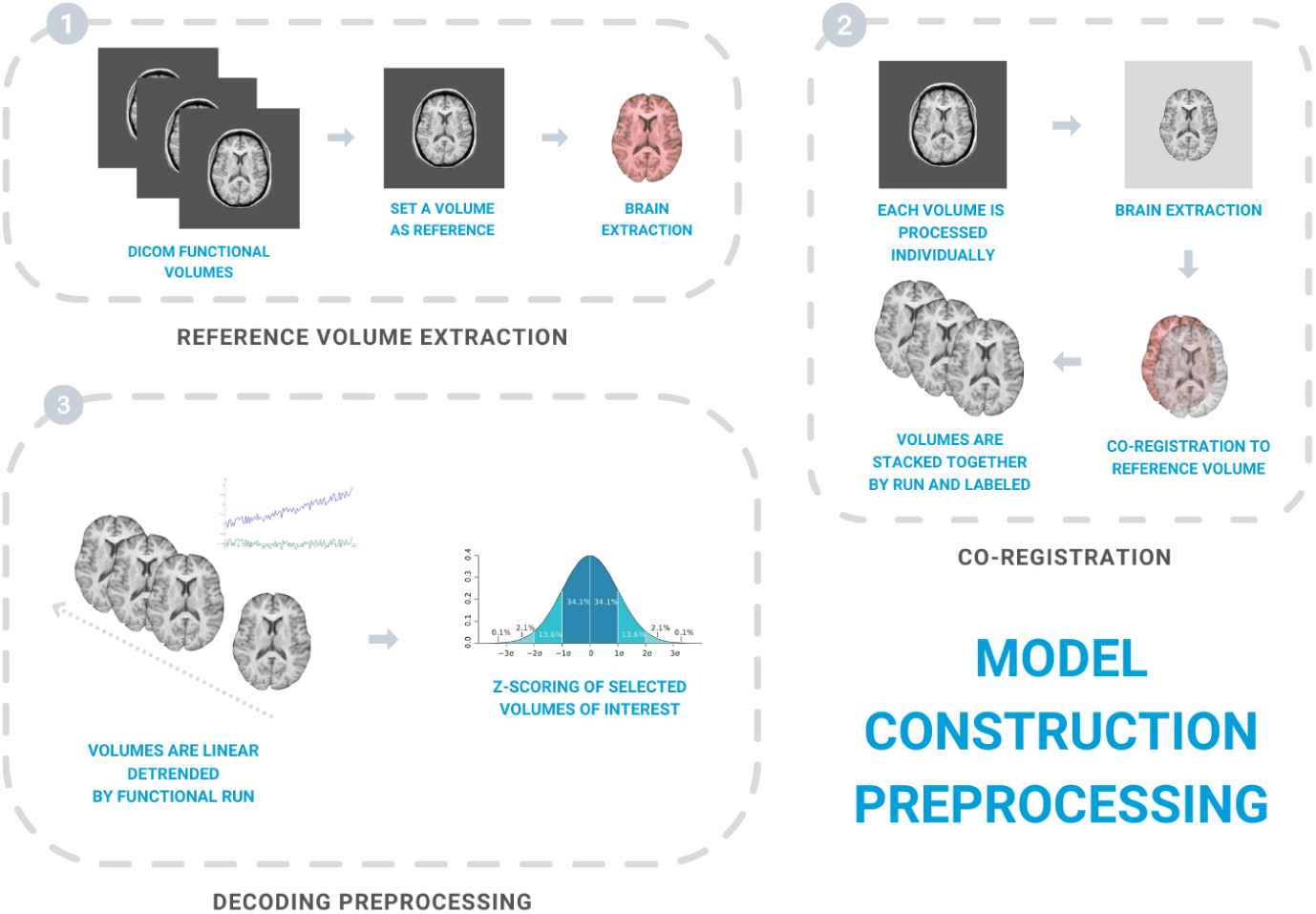
Illustration of the online preprocessing approach.

The server side is the main component and performs all the pre-processing and decoding operations. The client side is mostly used for providing stimulus and visual feedback to the participants in the MRI scanner, as well as for receiving the participants responses that are eventually communicated to the server side.

### System requirements for the server computer

For the server’s role in collecting and processing fMRI volumes in real-time, it is essential to have a computer with acceptable performance and ample storage capacity. The server computer must be capable of preprocessing and decoding volumes in less time than the selected fMRI repetition time (TR). Ideally, the server computer should run a Linux system such as Debian/Ubuntu 16+ or Redhat/CentOS 7+ to enable the use of either Docker or Singularity for the FIRMM software and either the Samba or rsync protocols for DICOM transfer.

Successful implementation of PyDecNef real-time scripts has been demonstrated on a computer running the CentOS Linux 7 operating system with the following specifications:

Memory: 31 GB

Processor: Intel© Core™ i9-9900K CPU @ 3.60GHz × 16

Graphics: Intel© HD Graphics (Coffeelake 3×8 GT2)

GNOME: Version 3.28.2

OS type: 64-bit

Disk: 2.0TB

#### FIRMM

To receive fMRI volumes from the MRI scanner host computer, it is necessary to install FIRMM version 2.1 or a newer version on the server computer. This installation facilitates data transfer using either the Samba or rsync methods, depending on whether the MRI scanner is Siemens or GE, respectively.

#### Neuroimage Analysis Software

To enhance the preprocessing speed of fMRI volumes and leverage the versatility offered by the Python language, pyDecNef incorporates various specialized neuroimage analysis software written in C. These software tools are seamlessly integrated into a unified Python workflow through the use of the Nipype library.

<u>AFNI installation</u>
<u>FSL installation</u>
<u>dcm2niix installation</u>

#### Python tools

The pyDecNef scripts are designed to run on Python 3.6 or any newer version, minimizing the reliance on external libraries and maximizing compatibility across different Python versions.It is essential to have Python installed on both the server and client computers.

Python Dependencies: The pipeline was tested in both Linux and Mac using Anaconda, we also recommend setting up a new conda environment for the package to have a clean installation of the required packages. The required packages are:

- Pandas
- Nipype
- Nilearn
- scikit-learn
- Colorama

Additionally we used Opensesame 4.0.0 at the client side for stimuli presentation to the participants. We recommend using python version 3.11.7 since it provides the best compatibility with the scripts and Opensesame 4.0.0 (check out the Opensesame <u>documentation</u> for more details).

### Libraries for data preprocessing and decoding

To minimize processing times, our pyDecNef Python framework leverages the Nipype interface module to connect Python with neuroimage software written in computationally efficient programming languages such as C, including dcm2nii, and AFNI. Further, to prevent potential overlap during processing of fMRI volumes, pyDecNef includes control measures that allow for independent processing of volumes in separate threads running in parallel. The preprocessing framework was designed to maximize the similarity of the data preprocessing for the decoder construction and the real-time processing during DecNef.

### Usage and description

The repository of PyDecNef scripts can be accessed through here https://github.com/pydecnef/PydecNef. Below we describe their functionality separately for each component.

### Data collection

The initial step of recording the fMRI data only uses the Opensesame script “images_presentation.osexp” on the client side. This is used to display the stimulus to the participant. The script is under the path https://github.com/pydecnef/PydecNef/blob/main/2_client/1_training/ and all details and information on the Opensesame framework and its installation can be found through this link https://osdoc.cogsci.nl/. The log files in CSV format generated by the Opensesame script are needed for the training process since they include the relevant classes and participant’s responses. These files can be moved to the server side through running the “1.sending_behavior_logs.py” script at the same path. On the server side the “1.receive_behavior_logs.py” script under the path https://github.com/pydecnef/PydecNef/tree/main/1_server/1_model_training/1_scripts/1_arranging_data needs to be run simultaneously as well. (Note: The process of moving the log files in this package is still in the experimental phase and can encounter some issues, alternatively these files can be moved manually by the user.)

### Decoder Construction

PyDecNef includes an example of the decoder construction paradigm including raw data and data processing scripts. Scripts can be found here:

https://github.com/pydecnef/PydecNef/tree/main/1_server/1_model_training

Decoding neurofeedback experiments typically commence with this session. During an experimental session, data is gathered to train a multivariate machine learning classifier, which is subsequently applied in real-time neurofeedback training sessions with participants. It is crucial to note that decoding neurofeedback experiments require high-performance classifiers (i..e. with a decoding performance above 70% ROC-AUC) that are capable of discriminating the relevant brain representation. Decoder construction paradigms should be designed with this criterion in mind. Simply replace the provided data with your own, adjust the scripts to match your experimental settings, and you are ready to proceed.

Before we start the training procedure, we need to pre-process the data. We start with dividing the raw fMRI data according to the runs by running the script “2.divide_raw_fmri_data_to_runs.py” under the path https://github.com/pydecnef/PydecNef/tree/main/1_server/1_model_training/1_scripts/1_arranging_data. The data preprocessing phase is then as simple as running every script under the path https://github.com/pydecnef/PydecNef/tree/main/1_server/1_model_training/1_scripts/2_preprocessing in a numerical order. All the generated pre-processed files will be under the path: ‘1.Server/1.model_training/2.data/preprocessed’. We note that Region of Interest (ROIs) masks are used in the experiment, the relevant files must be placed under the path: ‘1.Server/1.model_training/2.data/rois_masks’.

### fMRI Scan Preprocessing

#### Preprocessing Pipeline

The scans used for decoder construction and decoded neurofeedback are preprocessed using the same pipeline to minimize potential sources of variability and enhance the real time classifier performance. As a starting point, a functional scan is designated as the reference volume for both model construction and neurofeedback training sessions. All functional volumes from the model construction session are individually co-registered to the reference volume. Each volume is labeled with information pertaining to the run, onset time, stimuli category, etc. After labeling, volumes undergo linear detrending within each fMRI run. Subsequently, volumes of interest falling within the HRF peak of each trial are selected for model construction. These are then subjected to voxel-level Z-score normalization.

Preprocessing scripts for decoder construction can be found at: https://github.com/pydecnef/PydecNef/tree/main/1_server/1_model_training/1_scripts/2_prep rocessing organized by their execution order.

PyDecNef also includes a placeholder folder for raw data in DICOM format following the data structure specified in the preprocessing scripts.Placing your own data in this path https://github.com/pydecnef/PydecNef/tree/main/1_server/1_model_training/2.data/raw and modifying the scripts to match your experimental settings will prepare your pipeline for use.

Below we describe each of the steps.

##### 1. Reference Volume Extraction

The first preprocessing step involves selecting a raw functional volume to serve as the reference image for all other volumes in the model construction session, as well as for neurofeedback training sessions. Typically, fMRI runs begin with a few volumes (5-10 volumes) for image stabilization, which are then discarded. In this pipeline, the first raw volume following the MRI scanner’s heat-up phase is selected as the reference image. This DICOM volume is converted to NIfTI format (without compression) using dcm2niix. It is also skull-stripped using AFNI to facilitate co-registration with subsequent volumes.

##### 2. Volumes Co-Registration

Raw volumes are converted to NIfTI format, skull-stripped, and co-registered to the reference volume using a rigid body (6-parameter) transformations via AFNI’s 3dvolreg program. A 7th-order polynomial interpolation method is applied as the default spatial interpolation method during co-registration. To improve co-registration accuracy between scans, one can manipulate the “erode” or “clfrac” parameters in the brain extraction preprocessing step. Adjusting these parameters erodes the brain mask inwards to exclude skull and tissue fragments, improving co-registration accuracy. Note that if you change these parameters, they should also be updated within the co-registration pipeline function used during real-time neurofeedback training for that subject (i.e., “corregistration_pipeline.py”).

##### 3. Decoding

After labeling and co-registration the volumes, we proceed with training the machine learning classifier. Training the decoder requires the use of the scripts under the path https://github.com/pydecnef/PydecNef/tree/main/1_server/1_model_training/1_scripts/3_braindecoder. The “1.training_and_evaluating_models_wholebrain.py” is used to train the decoder with the whole brain data and the “2.training_and_evaluating_models_masked.py” is used for the Region of Interest (ROI) based masks applied to the whole-brain data. These scripts offer an optional arguments to choose the type of decoder. The default decoder is the extratrees classifier from the scikitlearn python library. The list of classifiers is: [“svm”, “svmlinear”, “decisiontree”, “extratree”, “randomforest”, “extratrees”, “bagging”, “gradientboosting”, “adaboost”, “naivebayes”, “kneighbors”, “mlp”, “sgd”, “logisticregression”], which can be accessed by adding a numerical argument to the script from 0 to 13 following the sequential order of appearance in the list. The argument 14 can be used to evaluate existing models; here the script will execute a leave one run out cross validation test. In order to do this the decoder’s file has to be named “evaluated_model” and placed under the path ‘1.Server/1.model_training/3.models/wholebrain’ for the whole brain decoder or placed under ‘1.Server/1.model_training/3.models/masked/{mask_name}’ with {mask_name} being the name of the mask used during the training of the decoder. Both scripts also incorporate some preprocessing steps that can be removed by changing the variable preprocessing to False inside the script. The script outputs a CSV file (info.csv) that includes the information about the model and the preprocessing pipeline.Using the PyDecNef decoder construction pipeline, we compared the different models on a single participant’s fMRI dataset from our previous study, which involved a perceptual classification task with dog and scissor images (see Margolles et al., 2024). The results are shown in Figure 3.

**Figure 3.**
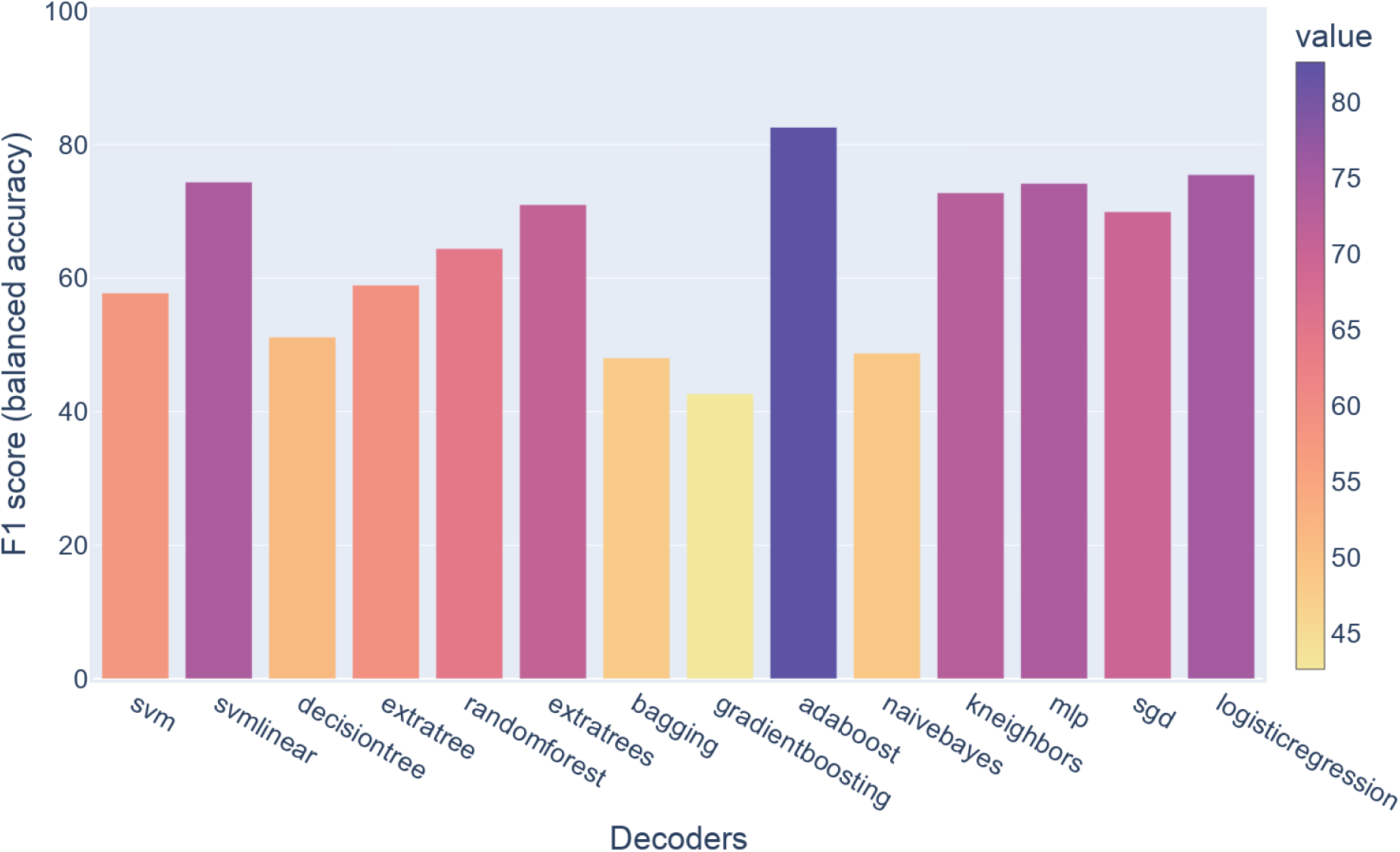
Illustration of the performance of different classifiers for an example participant.

### Real-time DecNef training

The neurofeedback phase is handled by the “2.neurofeedback” folders both in the server and the client side.

In the case of the server side, we need to provide the following required resources at the ‘1_server/2_neurofeedback/required_resources’:

- The ROI mask used for decoder training inside the path “1.server/2.neurofeedback/required_resources/{subject_tag}/masks” where {subject_tag} corresponds to the tag used for the participants, namely, in the form “sub-1”. If no mask was used in the decoder training phase, the mask is supposed to be example_func_deoblique_brainmask.nii that was generated inside the path: “1.server/1.model_training/2.data/preprocessed/example_func/” during the preprocessing phase.
- The decoder inside “1.server/2.neurofeedback/required_resources/{subject_tag}/models”
- The reference scan used for the volume realignment during the preprocessing needs to be placed inside the “training_session_ref_image” folder. This is the example_func_deoblique_brain.nii file generated during the preprocessing phase “1.model_training/1.scripts/2.preprocessing/1.generate_example_func.py” script.
- The training zscore data inside “training_zscoring_data” folder which include the two files, namely, zscoring_mean_array.npy and zscoring_std_array.npy generated in the training phase inside the “1.model_training/2.data/preprocessed/stacked_vols_of_interest/” folder. In the case that a mask is used this files are named zscoring_mean_array_{mask_name}.npy and zscoring_std_array_{mask_name}.npy in the “masked_stacked_vols_of_interest_{mask_name}” folder with {mask_name} is the name of the mask used in training.
- In the case of using co-adaptation, the training data and labels of the decoder need to be placed inside the “co_adaptation_base_training_stacked_vols_of_interest” folder. These files are detrended_zscored_stacked_vols_of_interest.nii.gz and detrended_zscored_stacked_vols_of_interest_labels.csv generated in the training phase inside the “1.model_training/2.data/preprocessed/stacked_vols_of_interest/” folder. Even in the case of using masks during the training process, these files are the same.

Additionally the config.ini file is the main tool to tweak the parameters and change the settings of the neurofeedback pipeline if needed before starting DecNef. After providing the required resources and checking the configuration, we can start the DecNef real-time training by running the https://github.com/pydecnef/PydecNef/blob/main/1_server/2_neurofeedback/1_realtime_fMRI/run_main.shs cript through the following command ‘sh run_main.sh’. The script will prompt you for the participant number, session number and run number. The script will break if there is some issues or problems with the parameters or data, for example, if you point to the wrong folder path regarding the incoming fMRI scans (i.e. this was left as /firmm/20240903.test in the config.ini file).

The run_main.sh script is designed to be the entry point for the scripts responsible for the real time fMRI decoded neurofeedback that can be found here: https://github.com/pydecnef/PydecNef/tree/main/1_server/2_neurofeedback/1_realtime_fMRI

This section includes all the Python scripts necessary for the server computer to preprocess and decode volumes during neurofeedback training sessions in real-time.

The main.py script is called by running the run_main.sh script, and integrates the “config,” “classes,” and “pipelines” modules together (https://github.com/pydecnef/PydecNef/tree/main/1_server/2_neurofeedback/1_realtime_fMRI/modules). This is the main script that has to be run from the server, followed by the stimulus presentation software, in order to run a real time fMRI experiment.

#### Config Module

exp_config.py: The primary experimental configuration file where you specify details such as the number of MRI scanner heat-up volumes, baseline volumes, hemodynamic response function peak limits, fMRI TRs, types of z-scoring procedures, decoding procedures, and essential files for neurofeedback decoding (e.g., regions of interest, machine learning models, reference volumes, z-scoring values).

connection_config.py: Facilitates the connection between scripts running on the server and client computers and enables data sharing between them.

listener.py: Matches specific client requests (e.g., from the experimental software) with custom server actions.

shared_instances.py: Enables the sharing of global variables and class objects instantiated in “main.py” across all modules.

#### Classes Module

classes.py: Defines attributes and methods for Python classes that interact to store information and process volumes in real-time. These include independent volumes, volume time series, independent trials, file watchers, client listeners, and data loggers.

#### Pipelines Module

corregistration_pipeline.py: This basic pipeline utilizes dcm2niix, AFNI, nipype, and nilearn to perform real-time corregistration of volumes to a reference volume from the model construction session. It also masks the volume using a region of interest mask file.

preproc_vol_to_timeseries_pipeline.py: Contains three different pipelines to detrend and z-score each neurofeedback training session volume using time series information from that fMRI session run.

trial_decoding_pipeline.py: Contains three different pipelines to decode neurofeedback training trials and volumes using a previously trained machine learning model.

Decoding_coadaptation.py: Performs decoder coadaptation for fMRI data. It adaptively updates a machine learning model (decoder) based on incoming neuroimaging volumes that meet specific criteria, allowing the model to generalize better across different data examples. The process involves loading existing models, evaluating new data points, and updating the model accordingly.

These scripts are intended to run in parallel with the **Python-based experimental software** (i.e. Opensesame) that controls stimuli, task and feedback presentation. These scripts can be found here: https://github.com/pydecnef/PydecNef/tree/main/2_client

On the client side, we need to run the Opensesame script that corresponds with our desired experimental task and stimuli. This script is responsible for presenting the stimulus and generating the feedback signal in the form of a disk (see Abdennour et al., 2025). This script will also generate log files for the different runs with details about the decoding accuracy and other important information about the real-time DecNef operation.

Note the connection_config.py file is needed to set the connection between server and client PCs. The scripts contain examples of static and dynamic neurofeedback paradigms built within the OpenSesame software (Mathôt, Schreij & Theeuwes, 2012). The static paradigm is the standard approach used in decoded neurofeedback involving the so-called “induction” runs in which the participant is required to self-regulate her brain activity in order to maximize the size of a feedback disk on each trial. Unbeknown to participants the size of the feedback is proportional to the probability of decoding the target state in the brain activity patterns. The dynamic paradigm involves the continuous decoding of every scan and hence could be used to “trigger” the presentation of a stimulus based on the decoded brain activity pattern (Debettencourt, Cohen, Lee, Norman. & Turk-Browne, 2015).. These two types of paradigms can be adapted for your specific experimental purposes.

#### fMRI Simulator (3.fMRI_simulator_realdata)

https://github.com/pydecnef/PydecNef/tree/main/1_server/2_neurofeedback/2_MRI_simulator

We also provide a simulation procedure to make sure everything is ready for the real experiment.

The simulation could be activated by changing the simulated_experiment variable to True in the config.ini file and including pre-recorded fMRI DICOM files in the path “1.server/2.neurofeedback/2.MRI_simulator/real_data”. Then, we need to run the 1.server/2.neurofeedback/2.MRI_simulator/generate.py script to simulate generating new DICOM files similar to the way we do with the MRI scanner, but before that we need to start by running the 1.realtime_fMRI/run_main.sh script and the Opensesame script in the same the way as the real-time DecNef procedure.

generate.py: Simulates an fMRI scanner working in real-time. This tool is used for offline testing of your decoded neurofeedback experimental paradigm, for instance, to ensure accurate synchronization between scripts in the server and client PCs, namely to simulate real time DecNef preprocessing and decoding (using raw volumes from a participant from the decoder construction session) alongside the stimulus and feedback presentation.

Beyond providing the building blocks for co-adaptive decoding, it is important to demonstrate that these components behave as expected in realistic DecNef settings. In the next section, we briefly summarise empirical benchmarks of the co-adaptive module based on offline simulations and a proof-of-concept real-time decoded neurofeedback experiment (Abdennour et al., 2025).

### Assessing the effect of co-adaptive training

To quantify the impact of co-adaptive training in PyDecNef, we leveraged the dataset from our previous DecNef study (Margolles et al., 2024) together with new real-time fMRI data. Full methodological details and statistical analyses are reported in Abdennour et al. (2025); here we provide a brief summary of the protocol and the main findings relevant for the validation of the PyDecNef co-adaptation tools.

In the offline simulations, a linear logistic regression decoder was first trained on perceptual data in which participants viewed images of dogs and scissors, labelled as living vs. non-living, within a bilateral fusiform mask. The resulting perceptual decoder was then applied to neurofeedback runs in which participants attempted to induce the target representation (living class) while receiving feedback based on the decoded probability of the target class. Co-adaptation was implemented using the Decoding_coadaptation.py module in PyDecNef: after each neurofeedback trial, fMRI volumes whose decoded probability for the target class exceeded a pre-defined confidence threshold (65% or 80% in separate analyses) were added to the training set, re-labelled as belonging to the target class, and the decoder was re-fit. At each co-adaptation step, the updated model was evaluated on the remaining neurofeedback volumes that did not meet the confidence threshold, using the F1-score as the main performance metric. To monitor potential drift away from the original target representation, we additionally tested the co-adapted decoder on left-out perceptual runs (leave-one-run-out cross-validation). Finally, to assess cross-domain generalisation, we compared the performance of the original and co-adapted decoders on independent mental imagery runs in which participants imagined the dogs and scissors stimuli (see Abdennour et al., 2025). These simulations showed that co-adaptation systematically improved decoding performance relative to the static perceptual decoder. When tested on neurofeedback data, the co-adapted decoder achieved higher F1-scores than the non-adaptive model, even when the amount of training data was matched, indicating that the performance gain could not be explained simply by increased sample size. Improvements were also observed when testing on the independent imagery runs, with the co-adapted decoder yielding more accurate discrimination between living and non-living imagery despite never being explicitly trained on imagery data. Importantly, the drift analysis revealed that the co-adapted decoder retained stable performance on left-out perceptual data across co-adaptation steps, suggesting that the incremental updates did not lead to overfitting or a loss of sensitivity to the original target representation.

Following the simulations, we evaluated the same co-adaptation procedure in a real-time DecNef experiment.. The model construction session comprised four perceptual runs identical to those described above. During the subsequent neurofeedback sessions, participants completed runs in which they attempted to induce either the living or non-living target representation while receiving disk-size feedback based on the decoded probability of the target class. Within each participant, we alternated between runs in which the decoder was kept fixed and runs in which co-adaptation was enabled, using the same confidence-based selection of neurofeedback volumes for incremental retraining.

Across participants, co-adaptive runs showed higher decoder performance than non-adaptive runs, indicating that the online updates facilitated participants’ ability to drive the system into the target state. Figure 4 illustrates this result. The effect was evident not only for the original living target but also when two participants returned for an additional session with the non-living (scissors) class as the target, where co-adaptation again led to improved decoding probabilities for the target representation. Taken together, these offline and real-time benchmarks demonstrate that the co-adaptation tools implemented in PyDecNef can be used to construct stable yet flexible decoders that progressively adapt to the statistics of neurofeedback data, thereby enhancing the effectiveness of DecNef protocols.

**Figure 4.**
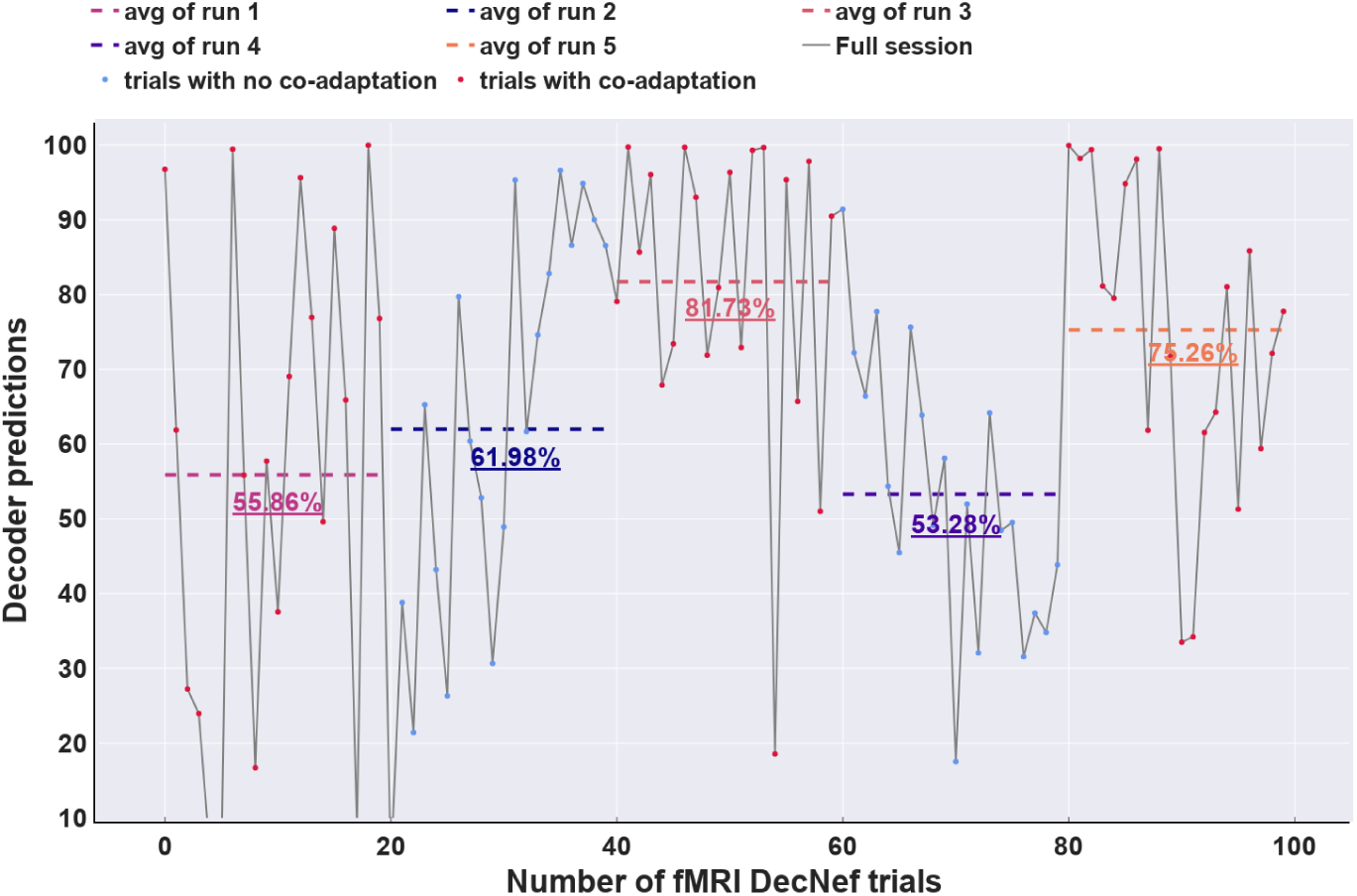
Illustration of decoder predictions for an example participant regarding the non-living class with co-adaptation (red) and without it (blue). Each of the fMRI trials in DecNef may have 2-3 examples.

### Comparison of different classification pipelines for DecNef training

The first release of PyDecNef (https://github.com/pedromargolles/pyDecNef) included a customised logistic regression classifier to improve DecNef performance. Previous studies on DecNef (Watanabe et al., 2017) employed classification models that often generated excessively binary predictions when applied to out-of-sample data. Consequently, these models hindered the utilization of nuanced neurofeedback signals to effectively facilitate participants’ learning in self-regulating their brain activity patterns. We developed a custom regression model in Tensorflow using Binary Cross-Entropy Loss to provide a more fine-grained decoding of the target class while accounting for noisy representations using soft-labeling (see https://github.com/pedromargolles/pyDecNef/tree/main/1.model_construction/sub-1/scripts/4.model_training). We note that this classifier does not currently support co-adaptation because retraining the model is too slow. For this reason, co-adaptation is not included in the current version of the package, although the classifier itself can still be used without it.

Furthermore, we standardized the fMRI scans during DecNef based on the data used for classifier training and further aimed to reduce the binomial nature of the predictions by averaging decoding probabilities from volumes within the HRF peak of each trial. The comparison of classification pipelines was performed based on the data from the DecNef study by Margolles,

Elosegi, Mei and Soto (2023). Data for decoder construction involved four perceptual runs and two mental imagery runs. The perceptual runs showed pictures of Dogs and Scissors. Following four perceptual runs, participants completed two mental imagery runs in which they were asked to vividly imagine one example picture from each of the visual categories.

We assessed how well our custom logistic regression classifier with BCE Loss performed compared to the standard and sparse logistic regression classifiers commonly used in DecNef research (Miyawaki et al., 2008; Yamashita et al., 2008). We evaluated the AUC and prediction probability distributions of the different decoders under three scenarios across participants: perception data, cross-domain generalization from perception to mental imagery data, and cross-domain generalization from perception to DecNef data.

We used the perception data to implement a leave-one-run-out cross-validation approach to estimate performance of the classifier. Here the decoder was iteratively trained using data from 3 perceptual runs, and then tested on the remaining run. For the standard logistic regression classifier, the maximum number of iterations was set to 5000. The L2 regularization penalty was applied to the logistic regression coefficients to prevent overfitting. The inverse of the regularization strength, known as C, was set to 1 to balance the model complexity and fit to the data. Lastly, the lbfgs solver was used to optimize the logistic regression objective function, since it is well-suited for small to medium-sized datasets. For the sparse logistic regression classifier (Yamashita et al., 2008), we used the Sparse Multinomial Logistic Regression (SMLR) Python implementation (Majima, 2015) available through this link: https://github.com/KamitaniLab/smlr. For comparability purposes we used default parameters. To be specific, we set a maximum number of iterations of 1000 for decoder training and used 1e-5 as the tolerance value of the stopping criteria. Visual noise trials were not used neither for training-testing standard nor sparse logistic regression classifiers. Finally, to maintain consistency with our experimental approach, we trained our custom logistic regression classifier with BCE Loss using the same parameters as mentioned in the Decoder training section. During the training process, the data was labeled with visual noise samples assigned to 0.5 during the training process. Training the decoder with Tensorflow required a validation set and accordingly we used a cross-validation approach in which the left-out run was divided into two halves to create a validation set and a test set. To obtain a more comprehensive assessment of the decoder performance, we repeated this cross-validation procedure by swapping the validation and test sets, ensuring that all data was used as a test set.

In the cross-domain generalization from perception to mental imagery data, the standard and sparse logistic regression classifiers were trained using all data from perception sessions (again, excluding visual noise trials) and then tested over the HRF peak volumes from the two mental imagery runs. The training data for our decoder with BCE loss consisted of volumes from ‘Dogs’, ‘Scissors’, and Visual noise trials from the first three perceptual runs. The validation set included ‘Dogs’ and ‘Scissors’ volumes from the 4th perceptual run, while the mental imagery data served as the test set.

Lastly, in cross-domain generalization from perception to DecNef data, the standard and sparse logistic regression classifiers were trained using ‘Dogs’ and ‘Scissors’ data from all perception runs and evaluated on all the HRF peak volumes within each DecNef training trial. To evaluate our custom logistic regression classifier, we utilized the pre-trained classifiers used in our experiment for each participant. We compared the different decoders by assessing the distribution of the predicted probabilities. Aggregated predicted probabilities for the three decoders across subjects can be visualized in Figure 5.

**Figure 5.**
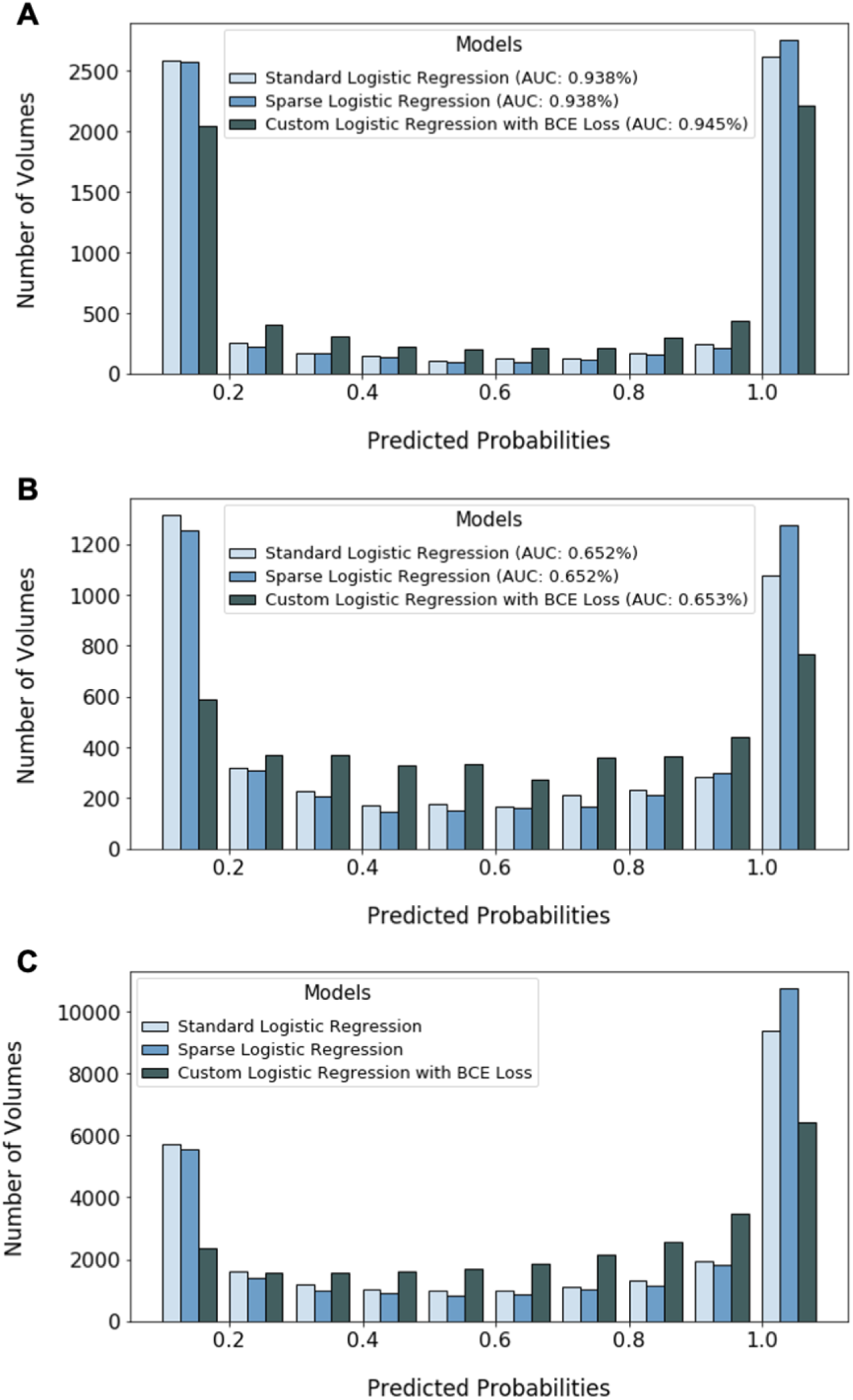
Histogram illustrating the predicted probability distribution (i.e., the number of volumes across participants) for various decoders, including standard logistic regression, sparse logistic regression, and our custom logistic regression classifier with Binary Cross-Entropy (BCE) Loss. This distribution is shown across different classification problems. (A) Perception: aggregated probabilities across the four cross-validation folds. (B) Visual imagery: aggregated cross-domain generalization probabilities based on a classifier trained on perception. (C) Aggregated cross-domain generalization probabilities during DecNef based on the decoder trained on perception and validated on imagery.

We used the Kolmogorov-Smirnov test to compare the distributions of predictions of the different decoders, which include standard vs. sparse logistic regression, standard vs. custom logistic regression, and sparse vs. custom logistic regression. In all scenarios, the decoder distributions were found to be significantly different from each other.

In the perception condition [Standard vs. Sparse - KS statistic: .056, p < .001; Standard vs. Custom - KS statistic: .149, p < .001; Sparse vs. Custom - KS statistic: .19, p < .001]; from perception to mental imagery [Standard vs. Sparse - KS statistic: .053, p < .001; Standard vs. Custom - KS statistic: .175, p < .001; Sparse vs. Custom - KS statistic: .161, p < .001]; and from perception to neurofeedback [Standard vs. Sparse - KS statistic: .059, p < .001; Standard vs. Custom - KS statistic: .156, p < .001; Sparse vs. Custom - KS statistic: .212, p < .001].

The probability distributions generated by the standard and sparse logistic regression classifiers exhibited overconfidence and greater similarity. These characteristics are particularly noticeable when examining cross-domain generalization scenarios. While maintaining performance, our customized classifier helped to mitigate the overly confident predictions in all scenarios.

### Assessing the influence of standardization methods on decoding performance

We conducted an analysis across participants using a similar approach to Oblak et al. (2019). This analysis leveraged the mental imagery data for decoder construction at our bilateral fusiform ROI (Margolles, Elosegi, Mei and Soto, 2023). Note that assessing the impact of normalization methods on the actual DecNef training data is difficult due to the absence of ground truth labels.

We evaluated five approaches: (1) no standardization, (2) standardization using the available run time series to each decoded volume, (3) standardization to the first 20 volumes of each mental imagery run, (4) standardization of decoded volumes based on the mean and standard deviation of the perceptual data, and (5) standardization using all the data within a run (as offline z-scoring method) (Figure 6A).

**Figure 6.**
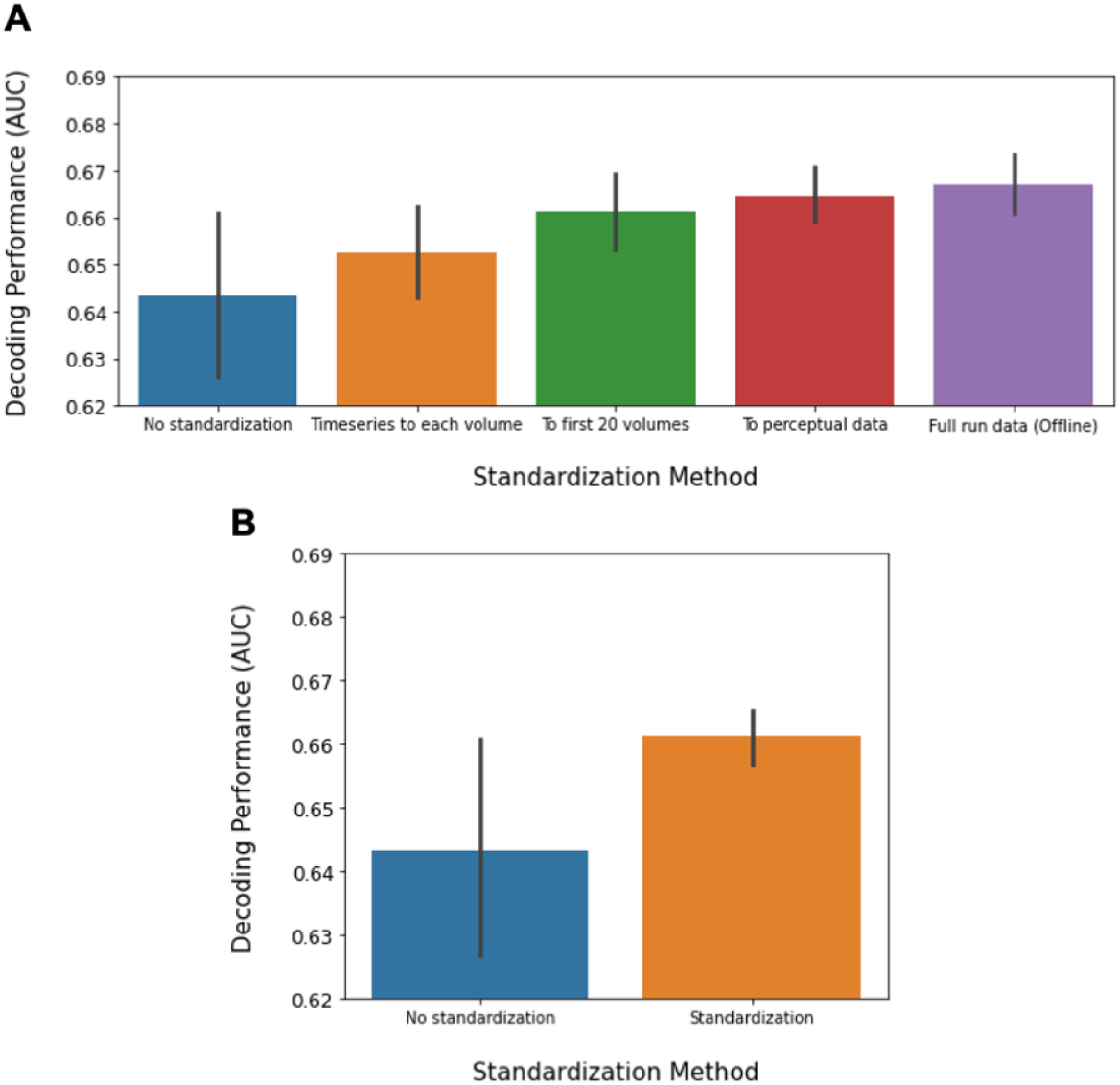
(A) Assessing the impact of various z-scoring techniques on cross-domain generalization from perception to mental imagery within our bilateral fusiform ROI across participants. In sequential order: 1 No standardization, 2 — Standardization using the available run time series to each decoded volume, 3 — Standardization to the initial 20 volumes of mental imagery runs, 4 — Standardization based on mean and standard deviation of perceptual data, and 5 — Standardization using all data within a run (offline z-scoring method). (B) Comparison of decoding performance with and without standardization, where standardization represents the averaged performance of the tested methods. Error bars represent 95% CIs according to the Cousineau–Morey method.

Even though our DecNef standardization approach, utilizing the mean and standard deviation of the training data for the decoder (i.e, perceptual data), demonstrated superior numerical decoding performance, in line with Oblak et al. (Oblak et al., 2019), a repeated measures ANOVA with Greenhouse-Geisser sphericity correction found no significant differences between the various z-scoring techniques [F(1.833, 31.165) = 2.592, p = .095, η p 2 = .132]. However, in contrast to their results, a paired t-test comparing the non-standardization and standardization of the data revealed no statistically significant differences [t = -1.588, p = .065] (Figure 6B).

## Discussion

PyDecNef provides an open, flexible and fully documented framework for implementing decoded neurofeedback protocols with real-time fMRI. By integrating data transfer, online preprocessing, decoder construction, and feedback presentation into a single modular platform, PyDecNef lowers the technical barrier for laboratories that wish to adopt DecNef and facilitates the sharing and replication of complex real-time pipelines across sites. In this way, PyDecNef contributes to a more transparent and collaborative ecosystem for closed-loop fMRI research.

Beyond providing basic infrastructure, PyDecNef incorporates specific methodological advances in decoder construction that are critical in the DecNef context. Our customised logistic regression models with soft-labels reduce overly binomial prediction profiles while preserving classification performance across perceptual, imagery and neurofeedback data. These properties translated into reliable learning curves in our recent DecNef study (Margolles et al., 2023), where participants progressively improved their ability to induce the targeted representation, indicating that the pipeline can support robust modulation of brain patterns during training.

A further development, now natively supported in the second version of PyDecNef, is co-adaptive training of the decoder. Using the same software components described here, Abdennour et al. (2025) showed in simulations and real-time experiments that incrementally updating the decoder with high-confidence neurofeedback examples improves decoding performance and participants’ ability to induce the target state, while drift analyses indicate that sensitivity to the original target representation is preserved. These findings suggest that co-adaptive DecNef can make training more personalised and efficient, potentially enhancing the magnitude and reliability of neural and behavioural effects and reducing the number of sessions required. Together, the static and co-adaptive tools implemented in PyDecNef offer a practical route towards more precise, adaptable and widely accessible decoded neurofeedback protocols.

## Open Science Statement

The scripts and documentation of the Python library for decoded neurofeedback (PyDecNef) can be found at: https://github.com/pydecnef/PydecNef. For the initial release of PyDecNef, see https://pedromargolles.github.io/pyDecNef/.

## Acknowledgements

P.M acknowledges support from an FPI grant from the Spanish Ministry of Economy and Competitiveness. P.E. acknowledges support from the Basque Government PREDOC grant. D.S. and N.A, acknowledge support from the IKUR programme and from the Basque Government through the BERC 2022-2025 program, from the Spanish Ministry of Economy and Competitiveness, through the ‘Severo Ochoa’ Programme for Centres/Units of Excellence (CEX2020-001010-S) and also from project grants PID2019-105494GB-I00 and PID2023-149267NB-I00.. The funders had no role in study design, data collection and analysis, decision to publish or preparation of the manuscript.

